# The cortical microtubules of *Toxoplasma gondii* underlie the helicity of parasite movement

**DOI:** 10.1101/2023.04.23.538011

**Authors:** Isadonna F. Tengganu, Luisa F. Arias Padilla, Jonathan Munera Lopez, Jun Liu, Peter T. Brown, John M. Murray, Ke Hu

**Author notes:** equal contribution (Isadonna F. Tengganu, Luisa F. Arias Padilla, Jonathan Munera Lopez).

## Abstract

Motility is essential for apicomplexan parasites to infect their hosts. In a three-dimensional (3-D) environment, the apicomplexan parasite *Toxoplasma gondii* moves along a helical path. The cortical microtubules, which are ultra-stable and spirally arranged, have been considered to be a structure that guides the long-distance movement of the parasite. Here we address the role of the cortical microtubules in parasite motility, invasion, and egress by utilizing a previously generated mutant (dubbed “TKO”) in which these microtubules are destabilized in mature parasites. We found that the cortical microtubules in ~ 80% of the non-dividing (i.e. daughter-free) TKO parasites are much shorter than normal. The extent of depolymerization is further exacerbated upon commencement of daughter formation or cold treatment, but parasite replication is not affected. In a 3-D Matrigel matrix, the TKO mutant moves directionally over long distances, but along trajectories significantly more linear (i.e. less helical) than those of wild-type parasites. Interestingly, this change in trajectory does not impact either movement speed in the matrix or the speed and behavior of the parasite’s entry into and egress from the host cell.

## INTRODUCTION

Apicomplexans are obligate intracellular parasites, many of which are important pathogens. For example, the malaria parasites, *Plasmodium* spp., kill more than half a million people each year (World-Health-Organization, 2022). *Toxoplasma gondii* permanently infects ~ 20% of the global human population and can infect any warm-blooded animal (Dubey, 2008). As obligate intracellular organisms, a successful infection requires the parasite to travel over a long distance through tissues with varying biochemical and biophysical properties (Hopp et al., 2015) as well as to enter into host cells. The latter requires a shorter travel distance, but perhaps more finesse, as the host cell plasma membrane needs to remain intact to allow the parasite to use host cell resources for proliferation.

*Toxoplasma gondii* moves along an irregular helical path in tissues and in a three-dimensional (3-D) matrix over tens of microns (Leung et al., 2014). Chemical inhibition as well as genetic manipulations showed that actin and a series of apical and cortical myosin motors provide the underlying force for the parasite motility (Dobrowolski et al., 1997; Gaskins et al., 2004; Graindorge et al., 2016; Heaslip et al., 2011; Meissner et al., 2002). The current model posits that the internal force is transmitted to the parasite surface through a coupling between the actomyosin complex and the secreted transmembrane adhesins (Opitz and Soldati, 2002). The characterization of these factors has significantly advanced the understanding of parasite motility (Dobrowolski et al., 1997; Frenal et al., 2010; Gaskins et al., 2004; Graindorge et al., 2016; Heaslip et al., 2011; Huynh and Carruthers, 2006; Huynh et al., 2003; Jacot et al., 2016; Meissner et al., 2002; Tosetti et al., 2019). However, while actin polymerization is critical for parasite movement, no native actin filaments have been detected, suggesting that the polymerization is likely to occur dynamically (Dobrowolski and Sibley, 1996; Mehta and Sibley, 2011; Periz et al., 2017; Sahoo et al., 2006; Schatten et al., 2003; Wetzel et al., 2003). Thus, it is still not known how the parasite is capable of traveling over long distances along a helical path. Intuitively, a stable, inherently helical cortical structure would be required for guiding continuous parasite movement over a long distance.

In *Toxoplasma,* one of the main cytoskeletal structures with a roughly helical arrangement is the set of cortex-associating (denoted as “subpellicular” or “cortical”) microtubules that are anchored at one end in a ring-like structure (the apical polar ring) at the parasite apex (Leung et al., 2017; Morrissette and Sibley, 2002a; Nichols and Chiappino, 1987). Among the apicomplexans, the microtubule cytoskeleton displays intriguing species and life-stage specific variations that coincide with substantial differences in physiology. The mature *Plasmodium* ookinete has an intricate cage of ~ 60 cortical microtubules (Bounkeua et al., 2010; Garnham et al., 1962; Wang et al., 2020), but the asexual blood-stage parasites only have a narrow band of 2-4 cortical microtubules that extends down one side of the parasite (Bannister and Mitchell, 1995; Morrissette and Sibley, 2002a). The *Toxoplasma* tachyzoite has consistently 22 cortical microtubules that are arranged in a spiral pattern. The parasite invests a considerable amount of resources in building and maintaining these polymers. Aside from the core building blocks, the tubulins, there is a suite of proteins that associate with the cortical microtubules in a region-specific manner (Leung et al., 2017; Liu et al., 2015; Liu et al., 2013; Tran et al., 2012). In mature parasites, these microtubules form an ultra-stable “rib cage” that extensively interacts with the inner membrane complex (IMC). They remain intact under treatments that depolymerize microtubules in mammalian cells within minutes (Hu et al., 2002; Morrissette et al., 1997). This extraordinary stability makes it difficult to assess the function of these polymers in the mature parasite.

Previously, a suite of proteins have been found to associate with the cortical microtubules. Some of them do not bind along the entire length of the microtubule, but are restricted to distinct regions (Liu et al., 2015; Liu et al., 2013; Tran et al., 2012). We discovered that, in mature parasites, TLAP2 and SPM1 together stabilize the portion of the microtubules distal to the apical section, which contains TLAP3 (Liu et al., 2015). In the absence of TLAP2 and SPM1, TLAP3 stabilizes the apical section of the cortical microtubules (Liu et al., 2015). Thus in the mutant that lacks TLAP2, SPM1 and TLAP3 (Δ*tlap2*Δ*spm1*Δ*tlap3,* denoted “TKO” for succinctness) the stability of the cortical microtubules in mature parasites is dramatically reduced. In the present work, we quantified the proportion of parasites with various levels of defects in the microtubule array under different conditions and found that ~ 80% of the TKO parasites that are not forming daughters (i.e. “non-dividing” in our terminology) have severely curtailed cortical microtubules. Depolymerization in mature TKO parasites becomes even more pronounced upon the initiation of daughter construction or cold treatment, the latter effect being partly reversible. While microtubule polymerization is essential for generating viable daughter cells, parasite replication is not affected by the destabilization of the cortical microtubules in the mature parasite. In a 3-D Matrigel matrix, the TKO mutant parasite moves as persistently as the wild-type parasite with similar velocity and net displacement. However, it travels along a path significantly more linear than the wild-type parasite. Interestingly, even though the cytolytic efficiency of the TKO parasite is significantly lower, live cell imaging revealed that the parasite’s behaviors during entry into and egress from the host cell are indistinguishable from those of the wild-type. This indicates that host cell entry and egress, which involve short-distance travel, is less sensitive to structural changes in the parasite than overall infection efficiency.

## RESULTS

### In TKO parasites, the daughter microtubule array is intact and replication proceeds normally

Previously, we showed qualitatively that the cortical microtubules in mature TKO parasites are destabilized compared with those in the wild-type parasites (Liu et al., 2015). Here, to quantify the extent of destabilization, we determined the proportions of TKO and wild-type parental (RHΔ*hx, (Donald and Roos, 1994)*) parasites that display various characteristics in the microtubule array. We first examined wild-type and TKO parasites that do not contain visible daughters based on anti-tubulin labeling (denoted as “non-dividing” in what follows). We found that while 100% of the wild-type parasites have a full, regularly arranged array, the cortical microtubules are severely curtailed in 79% (±8.5% STDEV) of TKO parasites (“short” phenotype) and sparse in the rest (Fig 1A) (Three independent experiments, total 490 parasites analyzed for the TKO and 246 parasites analyzed for the wild-type). The “short” and “sparse” phenotypes are defined as follows: “short”-microtubules at approximately normal density extend to less than 1/3 of the parasite body length and very few microtubules are longer than ~ 1/2 of the parasite body length. “sparse”-more microtubules are longer than 1/2 of the parasite body length but there are obvious gaps in the array. The extent of depolymerization in mature TKO parasites is more pronounced during cell division (Fig 1B-C). The proportion of parasites with nearly fully depolymerized cortical microtubules (“absent” category) was ~ 42% at the initiation stage of daughter construction and increased to ~ 98% prior to daughter emergence from the mother parasite (Fig 1C). For wild-type parasites, the cortical microtubules of the mother parasite in the initiation and elongation stages of daughter construction are similar to that of the non-dividing stage (Fig 1B-C). However, ~ 24% of mature parasites at the pre-emergence stage and 100% during daughter emergence have short microtubules.

**Figure 1.**
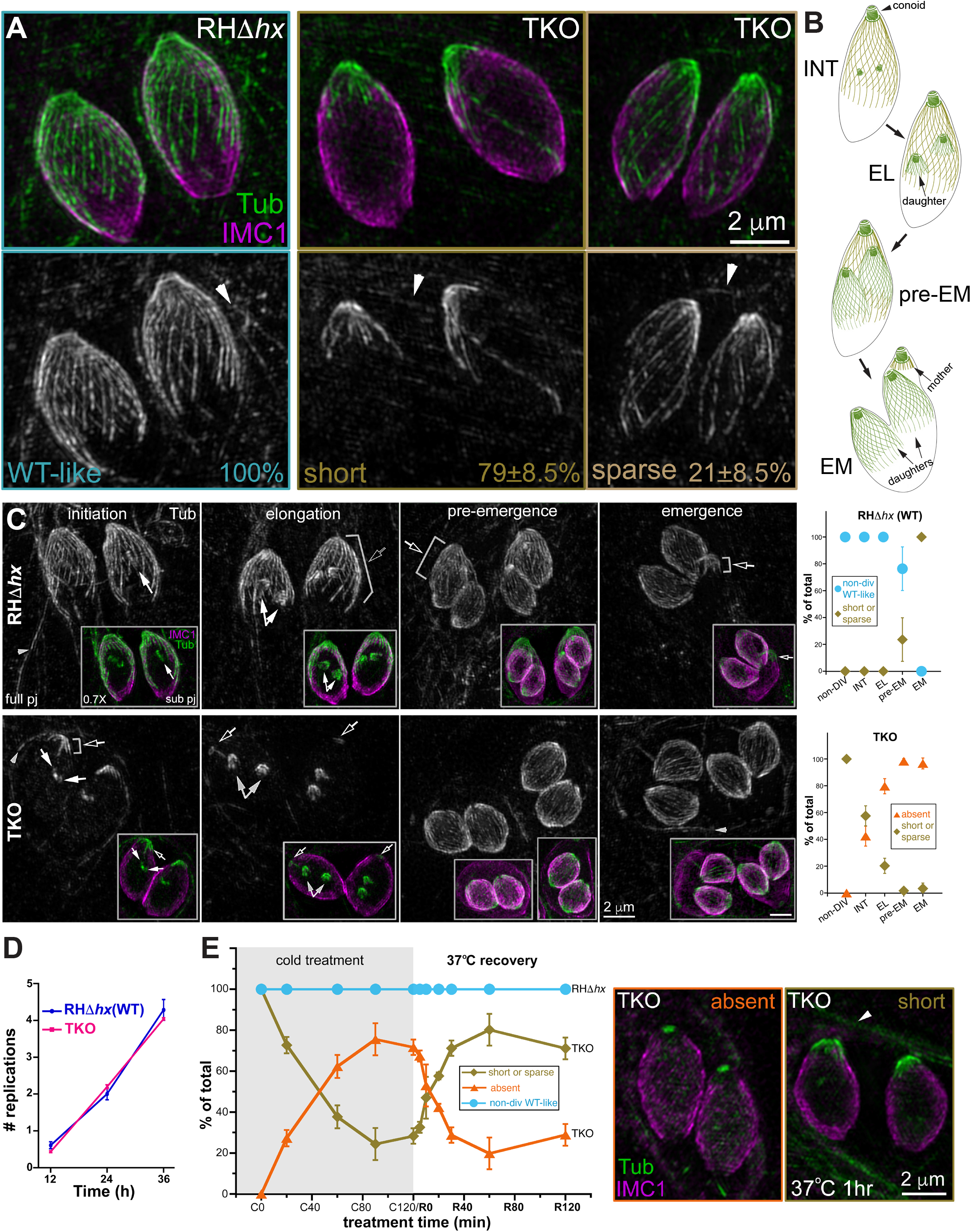
Cortical microtubules in intracellular mature TKO (*Δtlap2Δspm1Δtlap3*) and wild-type (RH*Δhx*) parasites. **A.** Projections of image stacks acquired by three-dimensional structured illumination microscopy (3D-SIM) of the non-dividing wild-type (RH*Δhx*) and TKO parasites. While 100% of the wild-type parasites have the typical spirally arranged full microtubule array, ~79% (±8.5% STDEV) of the TKO parasites have significantly curtailed/short microtubules. The cortical microtubules are sparse in the other 21%. Green and grayscale: tubulin labeling by a mouse anti-acetylated tubulin antibody, which also labels the microtubules in the host cell (arrowhead). Magenta: cortex labeling by a rabbit anti-IMC1 antibody. Arrowheads: host cell microtubules. **B.** Diagrams of wild-type parasites at the initiation (INT), elongation (EL), pre-emergence (pre-EM), and emergence (EM) stages of daughter construction. Cortical microtubules and the conoid are shown for the mother and daughter parasites. **C.** *Left*: Projections of 3D-SIM images of TKO and wild-type parasites at the initiation, elongation, pre-emergence, and emergence stage of daughter construction. Hollow arrows: cortical microtubules in the mother parasite. Arrows: daughter parasites. Arrowheads: host cell microtubules. Insets (0.7X) are projections that include a subset of the image stack (sub pj) to better display the structure of the daughter parasites, which are obscured in the full projections (full pj) for the wild-type parasites due to the cortical microtubules in the mother. Note that for the TKO parasite, due to the shortness or complete absence of the maternal microtubules, the daughters are as fully visible in the full projection as in the subset projection. *Right*: Quantification of the proportions of wild-type (top) and TKO (bottom) parasites that display various characteristics in the maternal cortical microtubule array at different stages, including the non-dividing (non-DIV) stage or the initiation (INT), elongation (EL), pre-emergence (pre-EM), and emergence (EM) stages of daughter construction. For the TKO parasites, the two patterns observed are “short or sparse” (*i.e.* similar to non-dividing TKO parasites) and “absent”. For the wild-type parasites, the two patterns observed are “non-dividing wild-type-like” or “short or sparse”. Error bars: standard error from three independent experiments. Total number of parasites analyzed: 88, 68, 50, 36 of TKO and 52, 62, 54, 49 of wild-type at the initiation, elongation, pre-emergence, or emergence stage, respectively. The data for the non-dividing stage is the same as shown in Fig 1A. **D.** Average number of cell divisions at 12, 24 or 36 hrs after infection in four independent experiments for RH*Δhx* (WT) and TKO parasites. Error bars: standard deviation **E.** *Left*: Quantification of the percentage of non-dividing wild-type (RHΔ*hx*) and TKO parasites that display various characteristics in the array of the cortical microtubules after being placed at ~ 7°C for 0, 20, 60, 90 or 120 min (C0-C120) as well as after recovery at 37°C for 5, 10, 20, 30, 60, or 120 min after 120 min of ~ 7°C incubation (R5-R120). For the TKO parasites, the two patterns observed are “short or sparse” or “absent”. For wild-type parasites, the cortical microtubules remain similar to those in untreated parasites under all conditions tested here (after cold-treatment and cold-treatment plus 37°C recovery). Error bars: standard error from three independent experiments. *Middle*: projection of 3D-SIM images of TKO parasites in which cortical microtubules are absent. *Right*: projection of 3D-SIM images of TKO parasites after 1hr of 37°C (R60) recovery, in which short cortical microtubules at the apical end are seen. Arrowhead: host cell microtubules.

We examined the average number of rounds of replication at 12, 24, and 36 hours after infection and found that the rate of replication of the TKO parasite is very similar to that of the wild-type parasite (Fig 1D). At first glance, it is surprising that a mutant that has destabilized cortical microtubules does not have a replication defect, because microtubule polymerization is coupled with cortex formation, and thus is required for daughter growth (Hu, 2008; Hu et al., 2006; Morrissette and Sibley, 2002b; Stokkermans et al., 1996). However, as we observed previously (Liu et al., 2015), in the same cell in which the cortical microtubules of the mother TKO parasite are depolymerized, microtubules do polymerize in growing daughters, which develop normally. These results reveal a cell-stage dependent microtubule depolymerizing activity that acts on the cortical microtubules in mature parasites but not the polymerizing microtubules in growing daughters. The depolymerization of cortical microtubules in mature TKO parasites becomes apparent at a much earlier stage of daughter formation and is much more complete than in the wild-type parasites, likely because the former are already destabilized due to the lack of TLAP2, TLAP3, and SPM1.

In our previous work, we showed that incubation at ~ 8°C can further destabilize the cortical microtubules of the TKO parasite (Liu et al., 2015). Here we characterize the time course of cold-induced disassembly of cortical MTs in mature TKO parasites and found that, as expected, the extent of depolymerization increased with the time of cold-treatment (Fig 1E). After ~ 2 hr of cold-treatment (C120), cortical MTs in ~ 72% of the parasites are nearly completely depolymerized with no visible MTs (“absent” category) and only a small tubulin-positive “cap” at the apical end. On the other hand, the array of cortical microtubules in wild-type parasites is not affected by the cold treatment. The cold-induced depolymerization in the TKO parasites is partially reversible (Fig 1E). When TKO cultures cold-treated for 2 hr were returned to 37°C, the cortical MTs were restored significantly after 20 min of the temperature shift. Upon 1hr of 37°C incubation (R60), ~ 80% of the parasites showed patterns (“short or sparse”) similar to those of untreated parasites, with the microtubules concentrated at the apical end of the parasite (Fig 1E), suggesting a strong preference for regrowth from pre-existing nucleation sites.

The preceding experiments were carried out with intracellular parasites residing in host cells. As the difference between intra and extracellular environment may affect subcellular structures in the parasite, we next examined the microtubule array in extracellular wild-type parental (RHΔ*hx*) and TKO (Δ*tlap2*Δ*spm1*Δ*tlap3*) parasites. We found that the distribution of different array morphologies was similar to what was observed in intracellular mature parasites (Fig 2A&B). More than 94% of the parental parasites have a full, regularly arranged array with and without cold-treatment. In contrast, the cortical microtubules are short in ~ 69% (±4.4% SEM), sparse in ~ 23% and absent in ~7.5% of the untreated TKO parasites. The cortical MTs are absent in ~ 68% (±6.4% SEM) of the extracellular parasites harvested after ~ 2 hr of cold-treatment (C120). Confirming the synergistic effect of the associated proteins, the cortical microtubules in 75% (±4.5% SEM) of the single knockout mutant Δ*tlap2* have wild-type-like arrangement. In contrast, only ~ 4% of the double knockout mutant (Δ*tlap2*Δ*spm1*) has a wild-type-like microtubule array. The cortical microtubules are short in ~ 58% (±6.7% SEM), sparse in ~ 29% and absent in ~ 8.7% of this line.

**Figure 2.**
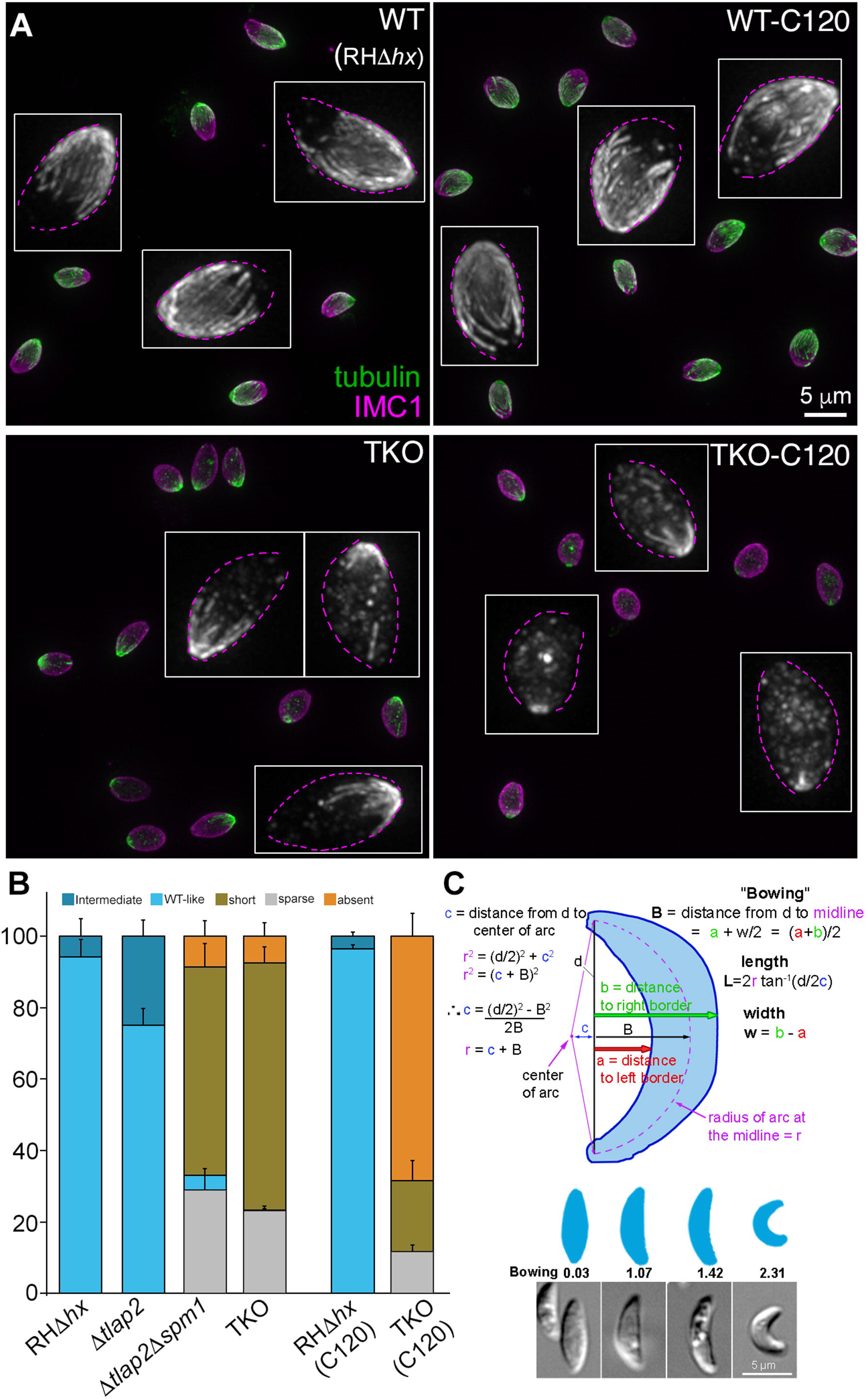
Comparison of cortical microtubule morphology and shape of extracellular wild-type parental (RH*Δhx*), *Δtlap2, Δtlap2Δspm1,* and TKO (*Δtlap2Δspm1Δtlap3*) parasites. **A.** Projections of image stacks of wide-field deconvolved images of extracellular RH*Δhx* and TKO (*Δtlap2Δspm1Δtlap3*) parasites with (C120) and without cold-treatment. Green and grayscale: tubulin labeling by a mouse anti-acetylated tubulin antibody. Magenta: cortex labeling by a rabbit anti-IMC1 antibody. Insets: Grayscale images of the tubulin labeling in a few parasites in each field displayed at 3X magnification. Dotted outlines were drawn based on IMC1 labeling. **B.** Quantification of the proportions of extracellular RH*Δhx*, *Δtlap2, Δtlap2Δspm1,* and TKO (*Δtlap2Δspm1Δtlap3*) parasites that display various characteristics in the cortical microtubule array. “Short”, “sparse”, and “absent” phenotypes are as defined in Figure 1. The “Intermediate” phenotype refers to arrays with detectable gaps but less severe than the “sparse” arrays. For RH*Δhx* and TKO parasites, measurements of parasites harvested after 2 hr (C120) of cold treatment are also included. Error bars: standard error. **C.** *Top:* Illustration for the calculation of length, width, and bowing of extracellular parasites. *Bottom*: diagrams and sample images of parasites that have different bowing indices. Quantification of length, width, and bowing of extracellular RH*Δhx*, *Δtlap2, Δtlap2Δspm1,* and TKO (*Δtlap2Δspm1Δtlap3*) parasites are included in Table 1.

**Table 1.** Quantification of length (L), width (W), length to width ratio (L/W), and Bowing (B) for four *T. gondii* strains. SEM: Standard error of the mean. P-values comparing the mutant strains (*Δtlap2, Δtlap2Δspm1,* and the TKO) with the RHΔ*hx* parent are from unpaired Student’s t-tests.

| Parasite strain<br>(Total number of<br>parasites analyzed) | L ($\mu m$) ± SEM<br>p value | W ($\mu m$) ± SEM<br>p value | L/W ± SEM<br>p value | B ($\mu m$) ± SEM<br>p value |
| --- | --- | --- | --- | --- |
| RH$\Delta hx$<br>(252) | 7.32± 0.06 | 2.05± 0.01 | 3.59± 0.03 | 1.42± 0.02 |
| $\Delta tlap2$<br>(207) | 7.09±0.04<br>**p = 0.002** | 1.91± 0.01<br>**p < 0.0001** | 3.74± 0.03<br>**p = 0.001** | 1.42± 0.02<br>p = 0.99 |
| $\Delta tlap2\Delta spm1$<br>(151) | 7.09± 0.04<br>**p = 0.004** | 2.02± 0.02<br>p = 0.20 | 3.55± 0.03<br>p = 0.39 | 1.37± 0.02<br>p = 0.10 |
| TKO<br>$\Delta tlap2\Delta spm1\Delta tlap3$<br>(181) | 7.04± 0.05<br>**p = 0.0007** | 2.14± 0.02<br>**p = 0.0003** | 3.33± 0.03<br>**p < 0.0001** | 1.07± 0.03<br>**p < 0.0001** |

Because of their stable association with the cortex, it has been hypothesized that the cortical microtubules support the characteristic crescent shape of *Toxoplasma.* We took advantage of the panel of mutants with varying degrees of defects in the microtubule cytoskeleton and examined their length, width, length-width ratio, and “bowing”. Length is defined as the distance from apical to basal tip measured along the arc that passes through both of them and through the center of the parasite, which is given by 2***r*** tan^-1^(***d***/2***C***) in Figure 2C. Bowing is a measurement related to curvature (see Figure 2C and Methods for definition). Because apparently straight parasites may be curved but viewed from the wrong direction, measurements were done on the most obviously bowed parasites for each line. The lengths of the microtubule mutants are slightly shorter than that of the wild-type parasite (Table 1). There is not a consistent correlation between microtubule defects and the width or length-width ratio of the parasite. The one parameter that stood out was the bowing index, which is significantly lower in the TKO parasite (1.07± 0.03 µm) than in all the other lines (1.37-1.42 µm) (Table 1). Interestingly, while the Δ*tlap2*Δ*spm1* parasite is much more similar in bowing to the RHΔ*hx* parental (1.37 vs 1.42 µm) than to TKO parasites, the structural defect of the microtubule array in the Δ*tlap2*Δ*spm1* is very pronounced and quite similar to that of the TKO (Fig 2B). This indicates that the effect of the structure of the microtubule array on parasite shape is likely to be nonlinear.

### The trajectory of movement of the TKO mutant is more linear than that of the wild-type parasite in 3-D Matrigel

In wild-type parasites, the stable, spirally arranged cortical microtubules have been proposed to guide directionality of parasite movement (Stadler et al., 2017) or to function as a spring that transmits energy to drive gliding motility (Pavlou et al., 2020). We tested these hypotheses by comparing the 3-D motile behaviors of the “wild-type” parent (RH*Δhx*) with those of the TKO parasite, taking advantage of the destabilized and curtailed microtubule array in the TKO. By modifying an established Matrigel-based 3-D motility assay (Leung et al., 2014), we obtained 3-D time-lapse images of the moving parasites under a label-free condition through Differential Interference Contrast (DIC) imaging. DIC imaging not only minimizes phototoxicity, but also enables the observation of parasite morphological changes as they move. However, while fluorescence images exhibit high-contrast that facilitates cell-tracking, automated tracking of cell positions in DIC image stacks is not as straightforward. We applied a series of edge detection filters, followed by erosion and dilation filters on the DIC images to derive images with contrast similar to fluorescence images (Fig 3A-B) that are amenable to 3-D cell tracking. Detection and tracking were carried out using the TrackMate plugin in Fiji, which generates the 3-D coordinates of the cell trajectories for further analysis and 3-D visualization (Fig 3C and Fig 4, Video S1-S4).

**Figure 3.**
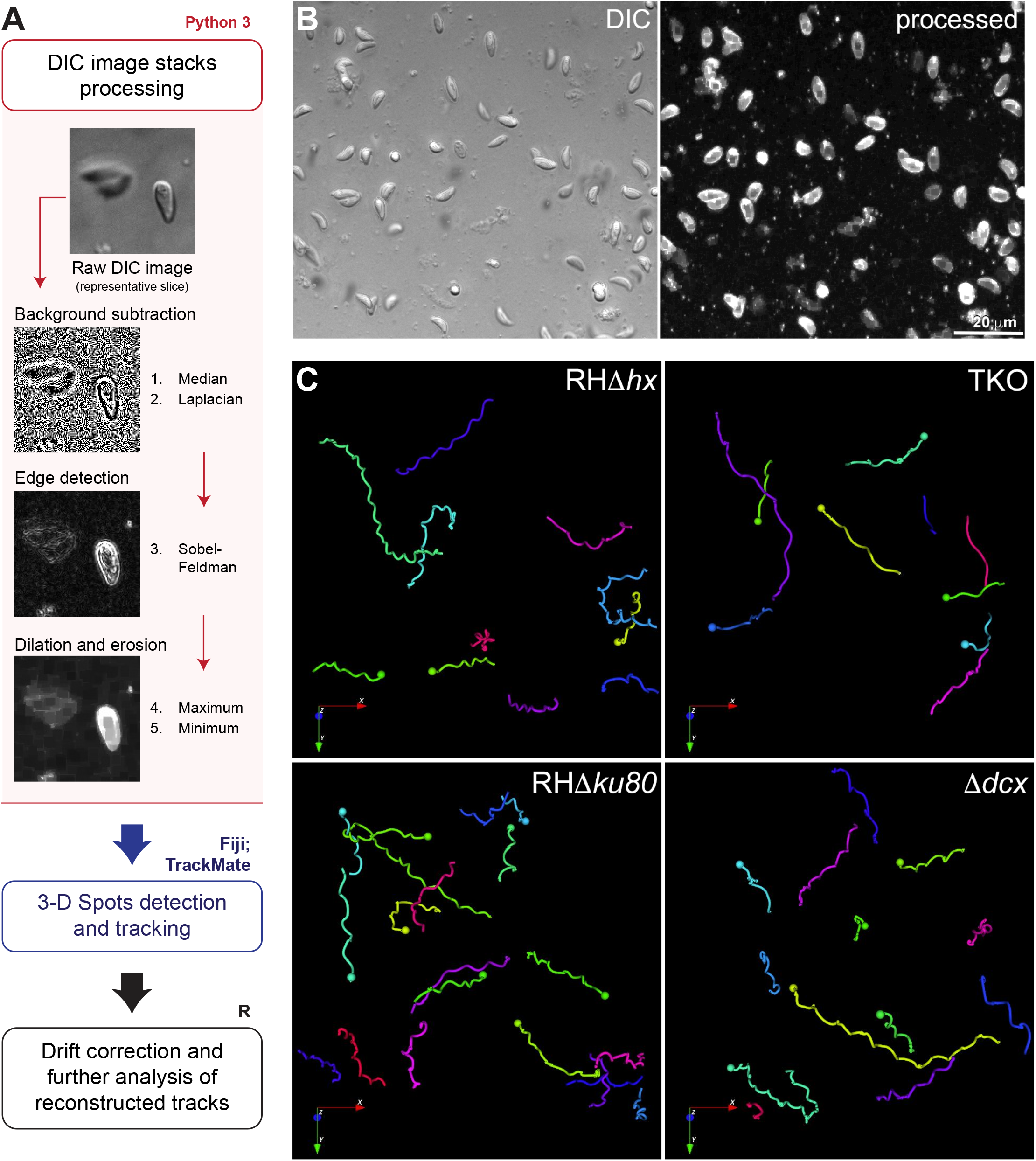
Workflow for image processing and automatic tracking of 3-D motility data acquired by DIC imaging. **A.** Schematic diagram of the image processing (pink outline and box), detection and tracking (blue outline), and tracks analysis (black outline) workflow for 3-D motility data acquired by DIC imaging. (pink) A representative area of a slice from a DIC image stack before processing and the resulting images corresponding to the filters applied on each processing step using Python3, showing the resulting high contrast image. For details, see Materials and Methods section. **B.** Focused composite image of a DIC image stack (41 slices; 40 µm) of extracellular wild-type parasites in Matrigel and corresponding processed image (represented by its maximum projection) (see also Video S1). **C.** Examples of 3-D tracks for motile RH*Δhx* parental, TKO, RH*Δku80* parental and *Δdcx* parasites reconstructed in the Icy software (see also Video S1-S4). Length of arrows = 16 μm.

**Figure 4.**
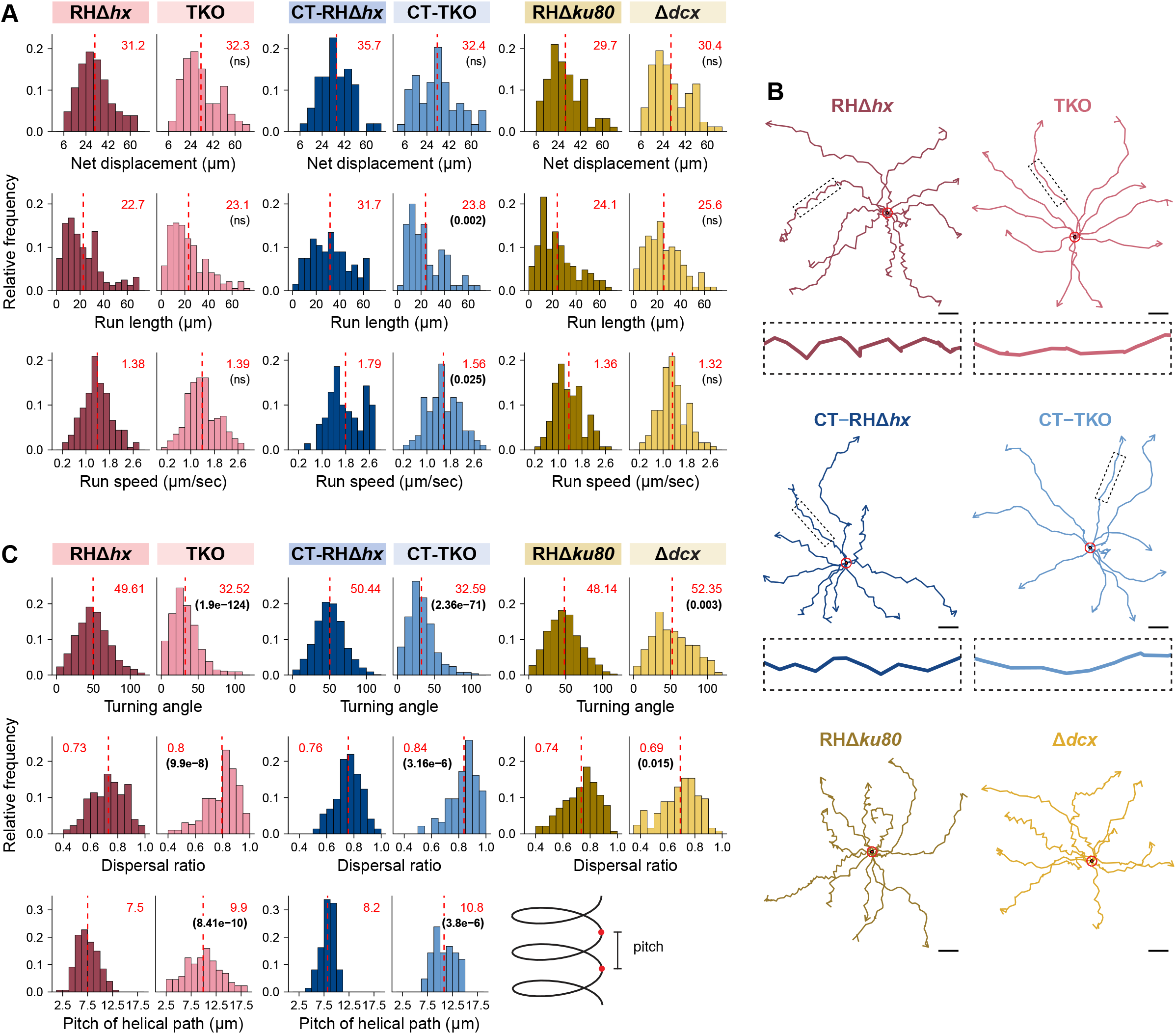
The 3-D trajectories of TKO parasites are more linear than those of parental parasites. **A.** Frequency distribution and statistical analysis of motion-related parameters including net displacement, run length (distance between pauses >5.4 sec), and run speed of RH*Δhx* parental and TKO without or with cold-treatment (CT), and RH*Δku80* parental and *Δdcx* parasites. For cold-treated TKO, intracellular parasite cultures were placed at 7 degree for two hours before harvesting. For the other samples/conditions, the parasites were harvested as described previously in (Munera Lopez et al., 2022). All motility assays were carried out at 37°C. **B.** Representative tracks projected on the (x, y) plane with superimposed origins (red circles) for RH*Δhx* parental and TKO without or with cold-treatment (CT), and RH*Δku80* parental and *Δdcx* parasites. Enlarged insets (4X) include the segments indicated by the dashed boxes in the trajectories. Scale bars: 10 µm. **C.** Frequency distribution and statistical analysis of helicity-related parameters including turning angle, dispersal ratio, and the pitch of RH*Δhx* parent and TKO, without or with cold-treatment (CT). An illustration defining pitch is included. Data for all parameters are also displayed for RH*Δku80* parental and *Δdcx* parasites except for the pitch. The mean values for individual histograms in A and C are shown at the top (in red) and also marked by the dashed line. P-values (in brackets under the mean values) were calculated by the Kruskal-Wallis test. ns: not significant.

To compare the impact of different helically arranged cytoskeletal structures, we also analyzed the motile behavior of a conoid mutant (*Δdcx,* (Nagayasu et al., 2016)) and its “wild-type” parent (RH*ΔhxΔku80*, abbreviated as “RH*Δku80*” (Fox et al., 2009; Huynh and Carruthers, 2009)) in parallel with those of TKO and its RH*Δhx* parent. The conoid is formed of helically arranged tubulin fibers (Hu et al., 2006; Hu et al., 2002; Nagayasu et al., 2016; Swedlow et al., 2002) (Fig 1B). In the *Δdcx* mutant, a structural component (TgDCX) was removed and the conoid structure becomes stunted and disorganized (Nagayasu et al., 2016). We found that on average, the motile percentage of the *Δdcx* is ~31% (486 parasites from 9 independent experiments analyzed), slightly lower than that of its “wild-type” parent (“RH*Δku80*”, 404 parasites from 8 independent experiments analyzed) (Table 2). The average net displacement, run-length (i.e., distance between pauses of >5.4 sec) and run speed (average speed during a run) of the Δ*dcx* parasite are similar to those of the wild-type parasites (30.4 vs 29.7 µm, ~25.6 vs 24.1 µm and 1.32 vs 1.36 µm/sec, respectively, Fig 4A). As previously reported (Leung et al., 2014), the wild-type parent moves along an irregular helical path. The average dispersal ratio (ratio of net displacement to total distance traveled of motile segments) of the trajectories is 0.74 (a value of 1 indicates a completely linear track). The average turning angle (relative angle of the parasite between two consecutive steps) is ~48°. The trajectories of the *Δdcx* are helical in nature (Fig 4B-C, Video S3-S4), with an average dispersal ratio of 0.69 and turning angle of ~ 52°, indicating slightly tighter turns.

**Table 2.** Fraction of motile parasites in the 3-D motility assay.

| Parasite strain | Mean percent motile ± SEM<br>(# of independent replicates) | Total number of parasites<br>analyzed |
| --- | --- | --- |
| RH$\Delta hx$ (TKO parent) | 40.2 ± 6.4 (N=6) | 413 |
| TKO<br>($\Delta tlap2\Delta spm1\Delta tlap3$) | 42.3 ± 6.8 (N=6) | 395 |
| RH$\Delta hx\Delta ku80$ ($\Delta dcx$ parent) | 36.5 ± 4.08 (N=8) | 404 |
| $\Delta dcx$ | 30.9 ± 5.0 (N=9) | 486 |

In contrast, the movement of the TKO parasite was visibly more linear than that of the wild-type parasite (Fig 4B, Video S1-S2). Confirming this visual impression, we found that the dispersal ratio of the TKO parasites is significantly higher than that of the wild-type parent (RH*Δhx*) (0.8 vs 0.73, p << 0.0001, Fig 4C). The turning angle of the TKO is significantly lower than the wild-type parasite (~32.5° vs ~ 50°, p << 0.0001). Both are consistent with the trajectories being more linear. For segments of trajectories during which the TKO parasite displays near helical movement, the average pitch of the helical turns is significantly longer than that of the wild-type parasite (9.9 µm vs 7.5 µm p << 0.0001). Other motility parameters are similar between the two lines. 40.2% of the RH*Δhx* parasites were motile (413 parasites from 6 independent experiments analyzed) (Table 2). ~42.3% of the TKO parasites were motile (395 parasites from 6 independent experiments analyzed). The average net displacement, run-length and run speed are not significantly different between the TKO and RH*Δhx* parasite (32.3 vs 31.2 µm, ~23.1 vs 22.7 µm and 1.39 vs 1.38 µm/sec, respectively, Fig 4A).

As shown in Figure 1A, there are some remnant cortical microtubules in untreated TKO parasites. To determine whether these remaining microtubules influence the helicity of parasite movement, we cold-treated (CT) the parasites for 2 hr, under which condition the cortical microtubules are nearly fully depolymerized in ~ 70% of the TKO parasites but those in the wild-type parasites are not affected (Fig 1E and Fig 2B). The parasites were then harvested and mixed with Matrigel for 3-D motility assay. The turning angle of the cold-treated TKO (CT-TKO) is nearly the same as that of the untreated TKO parasites (Fig 4B-C). As observed in untreated parasites, the cold-treated TKO parasite moves much more linearly than cold-treated wild-type parasite (CT-RHΔ*hx*), as indicated by the significant differences in turn angle (~32.6° vs ~ 50°), dispersal ratio (0.84 vs 0.76), and pitch (10.8 µm vs 8.2 µm).

To determine the impact of individual microtubule-associated proteins, we examined the 3D motility of *Δtlap2, Δtlap2Δspm1* along with the TKO and the wild-type parent (Fig 5). The loss of TLAP2 alone results in a mild defect in the microtubule array (Fig 2B) and a very modest change in the turning angle and dispersal ratio of the movement. The loss of TLAP2 and SPM1 together leads to pronounced defects in the microtubule array (Fig 2B) as well as significantly more linear movement as indicated by turning angle and dispersal ratio that are significantly different from the wild-type parent (Fig 5). The alteration in helicity of movement is therefore very much in line with the extent of perturbation of the microtubule array. We note that in this second set of 3D motility experiments, a new batch of Matrigel was used. It is well known that the mechanical properties of Matrigel can vary from batch to batch or change with prolonged storage (Aisenbrey and Murphy, 2020). In the second batch of Matrigel (Fig 5), the turning angles of both wild-type and TKO parasites are much lower than those of the corresponding lines measured earlier (Fig 4C). However, the differences in helicity (turning angle and dispersal ratio) remain highly significant between the mutant and the wild-type parent.

**Figure 5.**
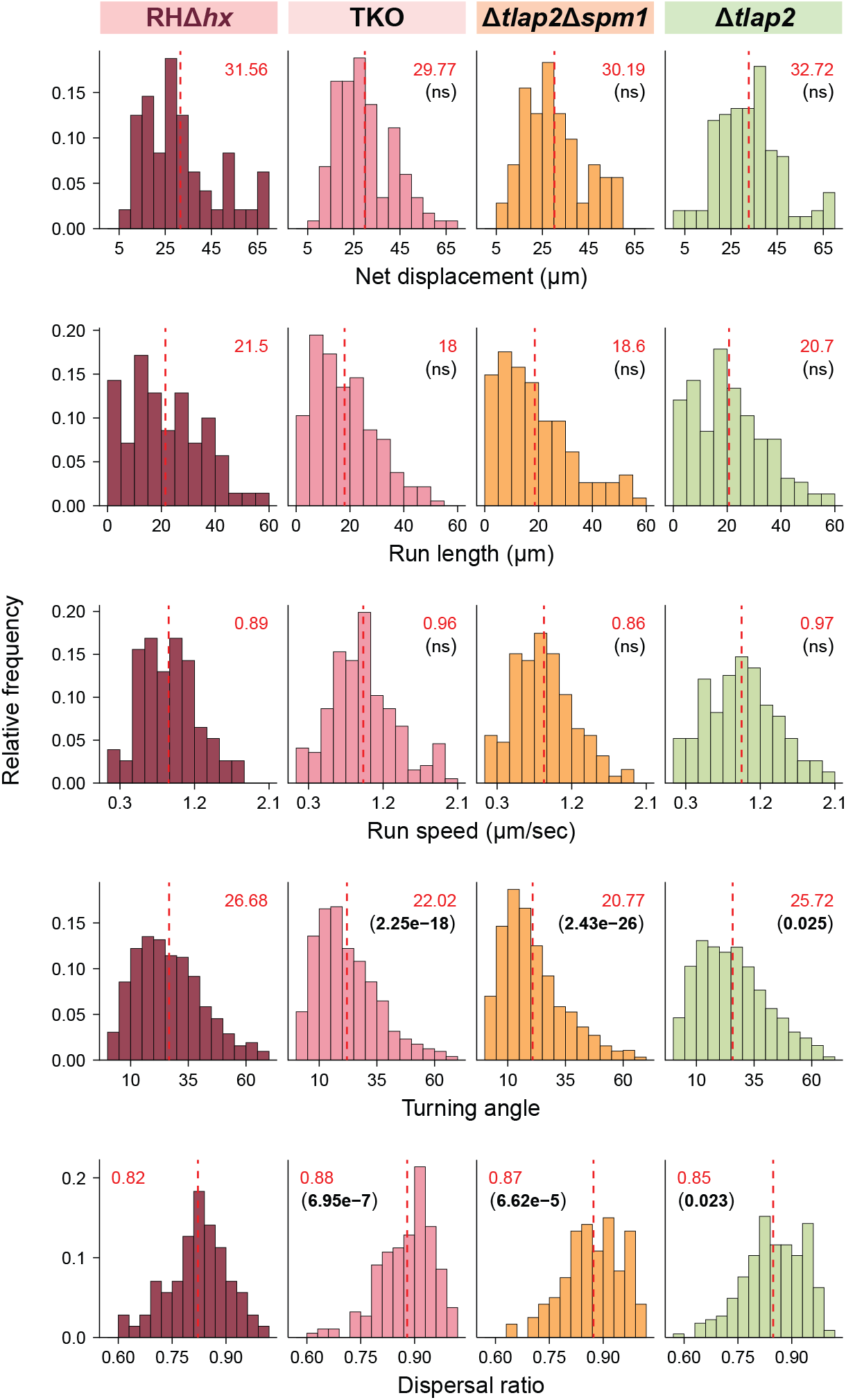
Comparison of 3-D motility of wild-type parental (RH*Δhx*), *Δtlap2, Δtlap2Δspm1,* and TKO (*Δtlap2Δspm1Δtlap3*) parasites. Frequency distribution and statistical analysis of net displacement, run length, run speed, turning angle and dispersal ratio of RH*Δhx*, *Δtlap2, Δtlap2Δspm1,* and TKO (*Δtlap2Δspm1Δtlap3*) parasites. Mean values for individual histograms are shown at the top (in red) and also marked by the dashed line. P-values (in brackets under the mean values) were calculated by the Kruskal-Wallis test.

### The apical concentration of micronemal vesicles is partially disrupted in the TKO parasite

Critical for effective parasite movement are transmembrane adhesins secreted from the membrane bound vesicles, micronemes (Carruthers et al., 2000; Carruthers and Sibley, 1997; Gras et al., 2017; Huynh and Carruthers, 2006; Rabenau et al., 2001). We have shown previously that micronemal vesicles co-align with the cortical microtubules (Leung et al., 2017). Here we quantify the different patterns of microneme distribution in TKO and its parent (RH*Δhx*). Micronemal vesicles were labeled by immunofluorescence using an antibody that recognizes MIC2, a major adhesin protein (Carruthers et al., 2000; Carruthers and Sibley, 1997; Gras et al., 2017; Huynh and Carruthers, 2006). The micronemal vesicles are enriched in an apical cap region (Fig 6A, ‘apical cap conc’) in 98% of the wild-type parasites, but only ~ 60% (±4.3% SEM) of the untreated TKO parasite (Fig 6A-B). When cold-treated for 2 hr (C120), this percentage further decreases to ~ 33% (±3.5% SEM) in the TKO parasites. The cold-treatment also has a modest impact on the wild-type parent, with parasites without the apical cap concentration increasing from ~ 2% to ~ 10% (Fig 6B). In parasites that lose the microneme enrichment at the apical cap, an intense concentration of MIC2 labeling remains at the parasite apex (Fig 6A, ‘no apical cap conc’). We note that a longitudinal bar is often observed at the parasite apex regardless of whether there is a prominent apical cap of microneme vesicles (Fig 6A, insets). This signal is probably from micronemal vesicles congregating in the conoid as observed by electron microscopy (Aquilini et al., 2021). To investigate whether the secretion of MIC2 is affected, we examined the amount of MIC2 secreted into the supernatant by extracellular RH*Δhx* parental and TKO parasites treated with the calcium ionophore A23187. We found that both lines respond robustly to the calcium ionophore treatment (Fig 6C and Fig S1). The TKO parasite on average secretes MIC2 at ~ 67% of the level of the RH*Δhx* parent (Fig S1), but the difference between the two lines is not statistically significant.

**Figure 6.**
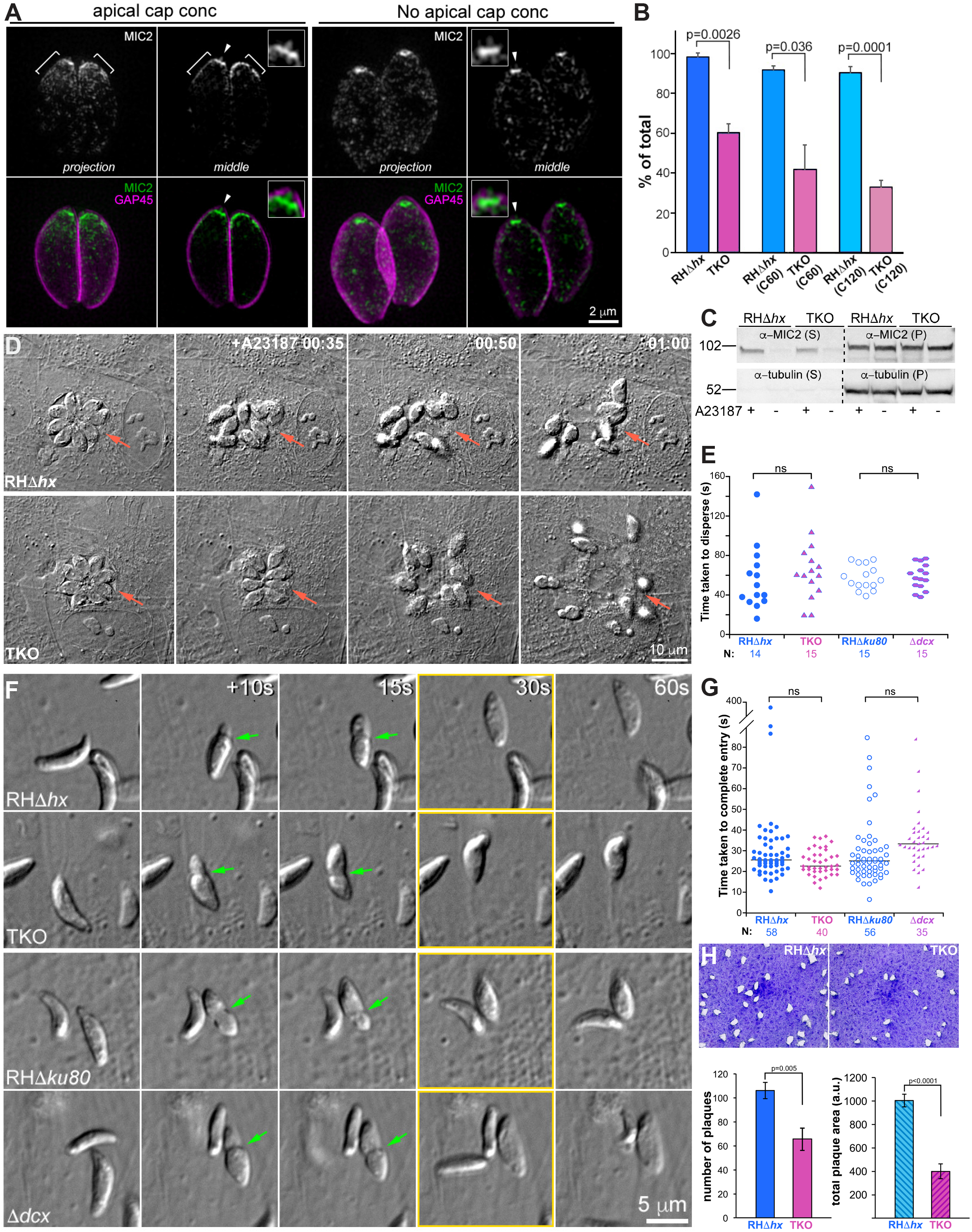
The speeds of entry and egress of TKO parasites are similar to those of parental parasites despite altered localization of micronemal vesicles, and lower invasion and plaquing efficiency. **A.** Projections and single sections of 3D-SIM images of representative intracellular parasites with and without an apical cap concentration of micronemal vesicles. Images of RH*Δhx* parasites and TKO parasites are used as examples for “apical cap concentration” and “no apical cap concentration”, respectively. Green and grayscale: microneme labeling by a mouse anti-MIC2 antibody. Magenta: cortex labeling by a rabbit anti-GAP45 antibody. Insets (shown at 2X, contrast enhanced to display weaker signals) include apex regions of parasites indicated by arrowheads. A longitudinal bar of MIC2 labeling at the apex is observed. **B.** Quantification of the percentage of RH*Δhx* parental and TKO parasites that have an apical concentration of micronemal vesicles. Three conditions are included: no treatment, and treatment at ~ 7°C for 60 (C60) or 120 min (C120) prior to the immunofluorescence labeling. P-values were calculated by unpaired Student’s t-tests. **C.** Western blots of the secreted (supernatant, S) and unsecreted (pellet, P) fractions of RH*Δhx* and TKO parasites with (+) or without (-) A23187 treatment. The blot was probed by antibodies against MIC2 and tubulin. The sizes of molecular weight markers (M) are indicated in kDa on the left. The contrast for images of the Western blot is enhanced and inverted to aid visualization of the bands. Please see Fig S1 for the full blot. **D.** Images selected from time-lapse experiments of intracellular RH*Δhx* parental and TKO parasites treated with 5 µM A23187 (also see Video S5). Parasitophorous vacuoles are indicated by the arrows. The RH*Δhx* parental and TKO parasite cultures were placed at ~ 7°C for 1 hr prior to the egress experiment, which was conducted at 37°C. **E.** Dot plots of time taken to disperse after treatment with 5 µM A23187 for RH*Δhx* parental, TKO, RH*Δku80* parental and *Δdcx* parasites. P-values were calculated by unpaired Student’s t-tests. ns: not significant. **F.** Images selected from time-lapse recording of RH*Δhx* parental, TKO, RH*Δku80* parental and *Δdcx* parasites in the process of invasion (also see Video S6). Frames marked in yellow: entry completed. Green arrows: constrictions formed during invasion. **G.** Dot plots of time taken for entry (i.e. time from parasite contact with the host cell at its apical end to completion of entry) for RH*Δhx* parental, TKO, RH*Δku80* parental and *Δdcx* parasites. P-values were calculated by unpaired Student’s t-tests. ns: not significant. **H.** *Top*: Plaques formed by the RH*Δhx* parental and TKO parasites. “plaques” are areas of the host cell culture lysed by repeated cycles of parasite invasion, replication, and egress. *Bottom*: Bar graphs that compare the number of plaques (left) and total plaque area (right) of the RH*Δhx* parental and TKO parasites. Five independent experiments were conducted. Error bars: standard error. P-values were calculated by unpaired Student’s t-tests.

### The infection efficiency of the TKO is lower than that of the wild-type parasite, while entry into and calcium-induced egress from the host cell is not affected

Parasite motility is critical for invasion into and egress from the host cell (Dobrowolski et al., 1997; Frenal et al., 2010; Gaskins et al., 2004; Graindorge et al., 2016; Heaslip et al., 2011; Jacot et al., 2016; Meissner et al., 2002; Munera Lopez et al., 2022; Tosetti et al., 2019). In addition, as the cortical microtubules are stable polymers that associate extensively with the parasite cortex, it is conceivable that they might provide mechanical support to the cortex when the parasites penetrate the membrane of the host cell during entry or exit. We therefore used video microscopy to determine whether the destabilization of the cortical microtubules in the TKO parasite might affect any of these processes. To examine parasite egress, the cultures infected by TKO or RH*Δhx* parental parasites were subjected to cold-treatment for 1 hr and then placed at 37°C for egress induced by A23187. Both TKO and wild-type parasites egressed actively and efficiently (Fig 6D, Video S5). Within 150 seconds, egress was completed in >93% of vacuoles for the wild-type parasites and in 100% of vacuoles for the TKO parasites. Of the egressed vacuoles, the average time taken to disperse is ~ 56 seconds for the wild-type parasite (n=14) and ~ 66 seconds for the TKO parasite (n=15) (p value not significant) (Fig 6E). In parallel, we also analyzed the egress behaviors of the *Δdcx* and its RH*Δku80* parent. We found that both lines egressed efficiently (Fig 6E), with an average disperse time of ~ 57.7 and ~ 58.3 seconds for RH*Δku80* (n=15) and *Δdcx* (n=15), respectively.

To assess parasite behavior during entry into the host cell, freshly harvested extracellular parasites were settled onto the host cell monolayer by a ~ 10 min pre-incubation on ice and then imaged by DIC at 37°C for 15 min (Fig 6F, Video S6). The number of entry events was recorded for four parasite lines: wild-type RHΔ*hx* parental, TKO, wild-type RHΔ*ku80* parental, and *Δdcx* parasite. We observed 58 events for RHΔ*hx* in 10 videos, 40 for TKO in 13 videos, 56 for RHΔ*ku80* in 9 videos, and 35 for *Δ*dcx in 22 videos. The median speed of host cell entry was 25.5 seconds for RHΔ*hx*, 22.5 seconds for the TKO, 25 seconds for the RHΔ*ku80* and 33 seconds for the *Δdcx* parasites (Fig 6G). We also did not observe any notable differences among these parasite lines in the morphological changes (*e.g.* the formation and resolution of constrictions) that accompany host cell entry.

The lower incidence of entry attempts observed in the time-lapse experiments for the TKO and Δ*dcx* indicates lower overall invasion efficiency. Indeed, *Δdcx* parasites were shown in our previous work (Nagayasu et al., 2016) to invade at a much lower efficiency than the wild-type parasites when invasion was assessed by a 2-color immunofluorescence assay that distinguishes invaded and uninvaded parasites based on different surface antigen accessibility (Carey et al., 2004; Mital et al., 2005). We determined the invasion efficiency of the TKO parasite using a similar approach and found that the TKO parasite invades at ~47 % of the level of its wild-type parent (p= 0.026, unpaired Student’s t-tests. # of parasites invaded/field: 25.2± 3.2 for RHΔ*hx* and 11.8± 1.1 for TKO). Consistent with lower efficiency in invasion, the TKO parasite generates ~ 60% of the number of plaques as the wild-type parasite (p ≈ 0.005) when examined by a plaque assay, where the parasites were incubated with the host cell monolayer unperturbed for 8 days (Fig 6H). The total area of the host cell monolayer lysed by TKO is 40% of the area lysed by the wild-type parasite (p < 0.0001), indicating significantly lower cytolytic efficiency. Thus, regarding the parasite’s ability to move helically or conoid structure, host cell entry is less sensitive than overall infection efficiency.

## DISCUSSION

The array of cortical microtubules is one of the major features of the cytoskeleton of *Toxoplasma* and other apicomplexan parasites. In *Toxoplasma,* the roots of the cortical microtubules are precisely located at evenly spaced intervals separated by cogwheel-like projections at the circumference of the apical polar ring (Leung et al., 2017; Morrissette and Sibley, 2002a; Nichols and Chiappino, 1987). The cortical microtubules are co-constructed and associated extensively with the inner membrane complex (IMC), guiding its formation during parasite replication (Hu et al., 2002; Morrissette et al., 1997). Inhibition of polymerization in replicating parasites blocks the elongation of daughter microtubules, and results in distorted daughter cortex and failed cell division (Hu, 2008; Hu et al., 2006; Morrissette and Sibley, 2002b; Stokkermans et al., 1996). In mature *Toxoplasma*, the cortical microtubules are extraordinarily stable and resistant to drug or cold-temperature treatments that readily depolymerize mammalian microtubules (Hu, 2008; Hu et al., 2006; Morrissette and Sibley, 2002b). It was proposed that the cortical microtubules might have various mechanical roles in facilitating parasite motility and invasion (Del Rosario et al., 2019; Pavlou et al., 2020; Stadler et al., 2017). However, the extraordinary stability of these polymers poses a major barrier against directly testing these hypotheses.

Previously, we generated a mutant (Δ*tlap2*Δ*spm1*Δ*tlap3,* a.k.a.“TKO”) in which the cortical microtubules are destabilized in the mature parasites while continuing to grow normally in developing daughters (Liu et al., 2015). Here, by quantifying the proportion of TKO parasites with different levels of defects in the array of cortical microtubules, we now have a more complete view of the extent of depolymerization in the non-dividing and dividing parasites. We found that 1) ~ 80% of untreated, non-dividing TKO parasites have severely shortened microtubules and 2) the extent of depolymerization in the mature parasite increases markedly when the daughters start to form, but the development of daughters is not affected. Examining cortical microtubules in the wild-type also revealed a small proportion of mature parasites with shortened cortical microtubules at the end of daughter construction. This suggests that microtubule depolymerization in the mother parasite might be a part of the normal preparation for disassembling maternal structures into component parts for possible recycling in the new generation. The cortical microtubules in mature TKO parasites are likely further sensitized to the depolymerizing activity due to the lack of TLAP2, SPM1, and TLAP3.

During division, the cortical microtubules of daughters grow continuously and do not display dynamic instability (Hu et al., 2002), indicating that these microtubules are stabilized by associated proteins. TLAP2, SPM1, and TLAP3 are recruited to the cortical microtubules of both mother and daughters (Liu et al., 2015; Tran et al., 2012) (Fig S2), yet they have differential effects on the stability of these two sets of microtubules sharing the same cytoplasm. In the same TKO parasite, while the maternal microtubules are destabilized, the daughter microtubules grow normally. This apparent specific stabilizing activity of TLAP2, SPM1, and TLAP3 for mature cortical microtubules could be because daughter cortical microtubules acquire additional specific stabilizing factors. Of course, for those (hypothetical) factors, we would need to answer how they differentially act on the cortical microtubules of the mother and developing daughters, which coexist in the same cytoplasm.

When incubated at low temperature, the cortical microtubules in the TKO parasite can be fully depolymerized (Liu et al., 2015). Here we show that this depolymerization is reversible: when the parasite was placed at 37°C after cold-treatment, short cortical microtubules were observed concentrated at the apical end of the parasite. This is consistent with the current hypothesis that the apical polar ring is the nucleating center for the cortical microtubules (Leung et al., 2017; Morrissette and Sibley, 2002a; Nichols and Chiappino, 1987). It is tempting to suggest that the nucleation complexes in the apical polar ring in the mature parasite remain active and are capable of re-nucleating new cortical microtubules after complete depolymerization. However, we cannot exclude the possibility that the new growth occurred from short microtubule stubs that remain associated with the apical polar ring, as a small cap of tubulin antibody labeling remains at the apical end even after prolonged cold treatment. If the latter is true, it suggests differential regulation of stability along the length of the cortical microtubules.

In the 3-D Matrigel, the TKO mutant moves much more linearly than the wild-type parasite, which travels along a helical path. The stable and spirally arranged cortical microtubules, with their extensive interactions with the cortex, could contribute to the helicity of the parasite movement by maintaining non-isotropic cortical properties (e.g. stiffness), and thus biasing the orientation at which the parasite interacts with its environment. In plant cells, the transmembrane cellulose synthesis complex is dynamically coupled to the cortical microtubules via cellulose synthases interacting (CSI1) protein (Bringmann et al., 2012). This results in the co-alignment of the extracellular cellulose fibers with the cortical microtubules on the other side of the plasma membrane. One could imagine that a similar mechanism might be involved in the confinement of the force-generating actomyosin apparatus associated with the inner membrane complex in *Toxoplasma*, although currently a CSI1-like coupler is unknown. The trajectory of the parasite movement might also be affected by the parasite’s shape. For example, in *Plasmodium berghei,* the knockout of IMC1h, which changes the shape of the ookinete, results in more linear movement (Kan et al., 2014). However, in the IMC1h *Plasmodium* mutant, while the cortical microtubules are still present and have a recognizable twist, they are also disorganized, which makes it difficult to disentangle the individual contributions from changes in parasite shape vs in the organization of the microtubules. We do not think the change in parasite shape is the likely explanation for the change in movement trajectory of the *Toxoplasma* microtubule mutants, because there does not appear to be a clear correlation between the parasite shape and linearity of movement. For example, the *Δtlap2Δspm1* parasite, which has a pronounced defect in the cortical microtubule array, moves much more linearly than the wild-type parasite even though their bowing indices are similar.

We note that while the net displacement, run length and run speed are similar in untreated parental and TKO parasites, these parameters are significantly higher for cold-treated wild-type parasites when compared with those for the cold-treated TKO (Fig 4A). This difference is due almost entirely to a response to the cold-treatment by the wild-type parasite, and the lack of response by the TKO. This lack of response by the TKO parasite indicates that the residual microtubules in the TKO under untreated conditions do not have a notable impact on parasite motility. The increase in speed and persistence of wild-type parasite movement after cold-treatment is not due to a direct effect of the microtubules on motility because 1) the cold-treatment does not have a detectable effect on the microtubule array in the wild-type parasite, and 2) the untreated wild-type parasite, which has a full complement and normal arrangement of cortical microtubules, has the same speed and persistence as the TKO parasite with few microtubules and a disordered array.

It has been proposed that the cortical microtubules act as a spring to store and release energy for helical gliding (Pavlou et al., 2020). However, our data suggest that these microtubules do not appear to act as a link in any energy relay vital for parasite movement. As discussed above, while the TKO mutant does not move helically, it is clearly capable of sustaining long-distance movement even under conditions where the TKO parasites were pre-treated with cold to nearly fully depolymerize the microtubules. Therefore the cortical microtubules do not play a role in maintaining directional long-distance movement. This means other higher-order structures are needed to limit the direction of force generation by the actomyosin complex. Whatever these structures might be, they need to be regularly arranged and stably associated with the parasite cortex. One candidate is the array of inner membrane particles (IMPs), which form periodic rows of IMPs that run the length along the longitudinal axis of the parasite (Morrissette et al., 1997). A subset of the IMP rows co-align with the cortical microtubules. The depletion of a potential IMP component (GAPM1a) in mature parasites results in decreased parasite displacement in the 3-D Matrigel as well as destabilized and disordered cortical microtubules (Harding et al., 2019). It is worth noting that, in contrast to the GAPM1a mutant, the net displacement of the TKO parasite is similar to that of the wild-type (Fig 4A). This separation of helicity from sustained parasite movement might be due to the cell-stage dependent nature of the stability of the cortical microtubules in the TKO parasite. The IMC cortex of the parasite is built concomitantly with the cortical microtubules during daughter formation. Because the daughter cortical microtubules grow normally in the TKO parasite, the structures in the IMC also likely form largely normally. Thus, we think that, in the mature TKO parasites, we are seeing mainly the impact of the microtubule destabilization itself, which manifests solely in the change from helical to linear movement.

Given the precision by which the cortical microtubules are constructed, the extensive suite of associated proteins (Leung et al., 2017; Liu et al., 2015; Liu et al., 2013; Tran et al., 2012), and their extraordinary stability, we had expected that microtubules would provide important mechanical support for the cortex in the mature parasite. To our surprise, the shape of the TKO parasite is still crescent in nature, and there is also no detectable difference in the behavior of the parasite during host cell entry and egress, when the parasite experiences considerable mechanical stress pushing through the host cell cortex. This indicates that other cortical structures, such as the inner membrane complex and associated intermediate filament-like network, are more important as structural support in mature parasites. Notably, the conoid structure mutant, Δ*dcx,* which invades with a much lower efficiency (Nagayasu et al., 2016), also showed wild-type like behavior during entry. This suggests that host cell entry is considerably less sensitive to the change in the structure of the cortical microtubules and conoid compared with extracellular migration during infection.

Lastly, TLAP2, SPM1, and TLAP3, the three proteins collectively responsible for the stability of cortical microtubules in mature *Toxoplasma*, have well-conserved orthologs in *Plasmodium* spp (Liu et al., 2015). It would be intriguing to determine whether the microtubule stabilizing function of these proteins, and the influence of cortical microtubule stability on parasite movement are also conserved among the apicomplexans. As the cortical microtubules in the developing daughter and the mature TKO parasites display distinct polymerization and depolymerization states, identifying the activities responsible for this maternal-filial distinction will generate insights into how the cortical microtubules of the mother and developing daughters (composed of the same polymer and less than half a micron apart) coexist in distinct biochemical states and how the switch from one state to the other occurs as the daughters become mature.

## MATERIALS AND METHODS

### T. gondii mutant strains and host cell cultures

Host cell and *T. gondii* tachyzoite parasite cultures were maintained as described previously (Leung et al., 2017; Liu et al., 2015; Munera Lopez et al., 2022; Roos et al., 1994). The TKO (Δ*tlap2*Δ*spm1*Δ*tlap3*), Δ*tlap2*Δ*spm1*, Δ*tlap2* mutants and the *mEmeraldFP-TLAP2* knock-in parasite were reported in (Liu et al., 2015). The Δ*dcx* parasite was reported in (Nagayasu et al., 2016).

### Temperature treatment of T. gondii for testing microtubule stability

Cultures of intracellular parasites growing in HFF cells plated in 3.5-cm glass-bottom dishes (MatTek Corporation, CAT# P35G-1.5-14-C or CAT# P35G-1.5-21-C) were incubated at 7°C for 20, 60, 90 or 120 min in L15 imaging media, which is Leibovitz’s L-15 (21083-027, Gibco-Life Technologies, Grand Island, NY) supplemented with 1% (vol/vol) heat-inactivated cosmic calf serum (CCS; SH30087.3; Hyclone, Logan, UT). For recovery, the cultures incubated at 7°C for 2 hours were subsequently placed at 37°C for 5, 10, 20, 30, 60 or 120 min in L15 imaging media. The samples were then processed for immunofluorescence as described below. Total number of intracellular parasites analyzed: 444, 422, 468, 423 of TKO and 246, 274, 300, 390 of wild-type cold-treated at 7°C for 20, 60, 90 or 120 min, respectively; 440, 428, 456, 496, 396, 360 of TKO, and 295, 168, 172, 166, 205, 230 of wild-type recovered at 37°C for 5, 10, 20, 30, 60, or 120 min, respectively.

### Immunofluorescence

All steps of the immunofluorescence labeling were performed at room temperature. For microtubule labeling of intracellular parasites (Fig 1), *T. gondii*–infected HFF monolayers growing in glass-bottom dishes were fixed in 3% (vol/vol) formaldehyde and 50% (vol/vol) methanol in 0.5X PBS for 15 min, permeabilized in 0.5% (vol/vol) Triton X-100 (TX-100) in PBS for 15 min, washed with PBS twice, followed by incubating with the primary antibody (mouse monoclonal anti-acetylated tubulin, 6-11B-1, 1:1000, Sigma T6793) and then with the secondary antibody (goat anti-mouse IgG-Alexa Fluor 488, A11029, Molecular Probes) at 1:1000, for 1 hr each. For labeling of the Inner Membrane Complex Protein 1 (IMC1), the samples were subsequently blocked with 1.5% BSA in PBS for 1 hour, then incubated for 1 hour with rabbit anti-IMC1 (a kind gift from Dr. Con Beckers, William Carey University) at 1:1000, and finally with goat anti-rabbit IgG-Alexa Fluor 568 (A11036, Molecular Probes) at 1:1000 for 1 hour. The antibodies were diluted in the blocking solution (i.e. 1.5% BSA in PBS).

For microtubule labeling of extracellular parasites (Fig 2), parasites were harvested as described in (Munera Lopez et al., 2022). To collect cold-treated parasites, cultures with intracellular parasites were placed at ~7°C for 2 hr as described above before harvesting. After harvesting, parasites were suspended in 3% (vol/vol) formaldehyde and 50% (vol/vol) methanol in 0.5X PBS and allowed to settle to the glass-bottom of the MatTek dishes for 15 min. The samples were then processed as described above for the intracellular parasites.

Immuofluorescence labeling of intracellular parasites with a mouse anti-TgMIC2 antibody, 6D10 (Carruthers et al., 2000) (a kind gift from Dr. Vern Carruthers, University of Michigan) was carried out as described in (Munera Lopez et al., 2022). A rabbit anti-GAP45 antibody (a kind gift from Dr Con Beckers, William Carey University) at 1:2000, followed by a goat anti-rabbit IgG-Alexa Fluor 488, (A11034, Molecular Probes) at 1:1000, was used to label the parasite cortex in these experiments (Fig 6A).

The labeling of the mEmeraldFP-TLAP2 knockin parasite with a mouse anti-ISP1 antibody (a kind gift from Dr. Peter Bradley, University of California, Los Angeles, (Beck et al., 2010)) (Fig S2) was carried out as described in (Liu et al., 2015).

### Three-dimensional structured-illumination microscopy

Three-dimensional structured-illumination microscopy (3D-SIM) image stacks were collected at z-increments of 0.125 µm using a DeltaVision OMX Flex imaging station (GE Healthcare-Applied Precision). A 60x oil immersion lens (numerical aperture [NA] 1.42) and immersion oil at refractive index 1.520 were used. Structured illumination image reconstructions and 3D projections were processed using the functions and software supplied by the manufacturer.

### Replication assay

The intracellular replication assay was performed as described in (Heaslip et al., 2011). Four independent experiments were performed. In each replicate, the number of parasites in ~ 100 vacuoles was counted for each strain at each time point.

### Shape analysis

Parasites were harvested from a heavily infected T12.5 flask of HFF by scraping, passing the suspension through a 23G needle, and filtering through a 3µm pore size filter. The suspension was centrifuged at 1500 x g for 3 min, the pellet was suspended 1 ml of L15 media containing 1% calf serum, and then centrifuged at 3500 x g for 2.5 min, discarding all but ~20 µL of the supernatant, which was used to resuspend the pellet. 10-15 µL of the concentrated parasite suspension was spread on a slide and covered with a coverslip. DIC images were collected as z-stacks of 7 to 19 slices in 0.8µm z-steps, using a 60x NA 1.3 silicon immersion lens. A custom script in the image processing software Semper (Saxton, 1996) was used to display the image, choose the best-focus z-layer, and pick parasites. The most obviously bowed parasites were selected for measurement because apparently straight parasites could be curved but viewed from the wrong direction. For each chosen parasite, shape parameters were calculated using the following manually identified features.

First, for each parasite image, a line was drawn from apical tip to basal tip. The coordinates of this line, with length denoted ***d***, were used to redisplay an enlarged image of the chosen parasite with the drawn line vertical, apical tip upwards. On this enlarged, re-oriented image, first the left, then the right-hand edge of the parasite was marked. Denote the distance from the apical-basal line to the left-hand border by ***a***, to the right-hand border by ***b***. Distances to the right of the line are positive, to the left are negative. The width of the parasite at its midpoint is given by ***b*** - ***a***, with ***a*** and ***b*** signed as defined. Length and bowing indices were then calculated from the coordinates of the marks as follows (see Figure 2C).

The “bowing”, defined as the distance, ***B***, from the drawn apical-basal line to the center (defined as half-way between the left-hand and right-hand edges) of the parasite is given by (***a*** + ***b***)/2, where ***a*** and ***b*** are signed numbers as defined above. Now consider an arc of an imaginary perfect circle that passes through the center of the parasite as well as through the endpoints of the previously drawn apical-basal line. Let ***C*** be the distance from the center of that circle to the midpoint of the apical-basal line, measured perpendicularly. Consider two radii, length ***r***, of that circle, one from the center of the circle to the apical tip of the parasite, the other from the center of the circle to the midline of the parasite. Referring to the diagram in Figure 2, it will be seen that for the first radius, ***r^2^*** = (***d***/2)^2^ + ***C^2^***, and for the second radius ***r^2^*** = (***B*** + ***C***)^2^. Solving for ***C*** gives ***C*** = [(***d***/2)^2^ - ***B***^2^]/2***B***, and ***r*** is then calculated as ***B*** + ***C*.**

If the parasite were not bowed at all, then the parasite length would be exactly ***d***.

We define the length (***L***) of the parasite to be the distance from apical to basal tip measured along the arc that passes through both of them and through the center of the parasite, which is given by 2***r*** tan^-1^(***d***/2***C***).

### 3-D motility assay and image acquisition

The 3-D motility assay for *Toxoplasma,* originally developed in (Leung et al., 2014), was carried out largely as described in (Munera Lopez et al., 2022). The environmental controller of a DeltaVision OMX Flex imaging station (GE Healthcare-Applied Precision) was set to heat the stage and objective lens at 37°C and allowed to equilibrate for >30 mins. A flow chamber was assembled as described in (Munera Lopez et al., 2022), then placed onto the heated stage at least 10 minutes before imaging. For cold-treated parasites, cultures with intracellular parasites were placed at ~ 7°C for 2 hr, before harvesting as described in (Munera Lopez et al., 2022). The harvested parasite suspension was kept on ice for ~30 sec, then mixed with an equal volume of Matrigel (Corning, 356237) on ice and pipetted into the pre-heated flow chamber. The chamber was then sealed with a polymerizable liquid silicone rubber (Nobilium Inc, Nobilsil #7453+7434) and image acquisition was started within ~2-3 mins after the suspension had entered the 37°C chamber.

For each imaging experiment, 151–181 image stacks were acquired at 1.36 sec intervals (7 msec exposure per optical section). Each stack consisted of 41 optical sections, spaced 1 µm apart in the Z-plane (40 µm stacks), with field of view of 1500×1500 pixels, binning 2×2, with a final dimension of 750 × 750 pixels. Image stacks were acquired from at least ~10 µm above the coverslip.

To represent the 3-D DIC images in 2-D format (Fig 3 and Video S1&3), we performed focus-stacking using Fiji by applying the Extended Depth of Field Sobel Projection plugin from (Forster et al., 2004; Schindelin et al., 2012)

### DIC image stacks processing and cell tracking

DIC images were processed using our custom scripts extended from (Munera Lopez et al., 2022). High contrast images were generated by subjecting the DIC images to 1) a 3-D median filter (kernel nx = ny = nz = 2 px) to reduce noise, 2) a 2-D Laplacian filter to attenuate out-of-focus portions of the cells, and 3) a 3-D gradient filter of Sobel type for edge detection. A dilation and erosion process using a maximum filter followed by a minimum filter (kernel nz = 3 px, nx = ny =13 px) was then applied. 2 pixels from all x and y edges were cropped before tracking to avoid edge artifacts. We implemented the imaging filters using Python 3 to enable GPU-acceleration using the CuPy plugin (Harris et al., 2020; Okuta et al., 2017; Van Rossum, 2009). The TrackMate plugin in Fiji was then used to detect and track the parasites in the processed image stacks (Ershov et al., 2022; Schindelin et al., 2012; Tinevez et al., 2017). The Difference-of-Gaussian (DoG) detector was employed for spot detection with a diameter of 7 µm. To avoid artifacts associated with edge-effects and out-of-focus signal, we excluded spots detected within the first and last 1 µm of the z-dimension and 3 µm of each of the x and y dimension. The Kalman filter tracker was used to reconstruct the cell tracks, and false reconstructions (i.e., “tracks” between debris or random noise) were filtered out based on quality, track length, and duration. The values for these parameters were determined empirically. The resulting tracks were manually verified and errors such as misdetections and gaps were corrected by adding or deleting individual spots. To refine spot positions, we used the TrackMate spot fitting feature to fit a 3-D Gaussian with fixed radius on each spot. After conducting a second round of manual verification, we then separated the motile parasite tracks from the non-motile tracks by applying another set of filters based on displacement and speed. The 3-D coordinates of both the non-motile and motile tracks were copied to different comma-separated value text files (.csv) for analysis in R version 4.1.2 in RStudio (RStudioTeam, 2020; Team, 2021).

### Trajectory analysis

We used the R package celltrackR v 1.1.0 and our own custom functions to process and analyze the parasite trajectories (RStudioTeam, 2020; Team, 2021; Wortel et al., 2021). The complete list of the rest of the R packages we used for our analysis is at the end of this section. First, the global drifts in the data were corrected according to the workflow developed by (Wortel et al., 2021) and we calculated the drift velocity from the corresponding set of non-motile tracks. Then, non-motile ends (the very start or very end) of the parasite trajectories were trimmed to eliminate biases introduced by the asynchronous nature of the parasite motility. After trimming, only tracks with more than four steps (i.e., five tracking points) were included in the subsequent analysis.

In Figures 4 and 5, the net displacement is a simple Euclidean distance between the first point and the last point of the trajectories. The run length is the total distance traveled between two “pauses”, which we defined as an event when the parasite stops moving for more than 5.4 sec (>= 4 frames). “stop moving” means that the speed of a step (sub-track of length 1) is lower than a threshold. For each data set, the threshold value was calculated by taking the average step-speed of the non-motile tracks. We removed the track points that correspond to pause events, introducing gaps. The motile segments were then obtained by splitting the tracks around those gaps. The run speed was obtained by dividing a motile segment’s total distance (run length) by its total duration. For Figures 4C and 5, the dispersal ratio was calculated by taking the net displacement of the motile segments and dividing it by the run length, where a value of 1 indicates a completely linear track.

For analyzing the turning angle (Fig 4C and Fig 5), any stops that were not filtered out before (i.e., stops that lasted less than 4 frames) and steps with displacement of less than 10% of the parasite body length (~ 0.75 µm) were cut out from the motile segments before splitting into sub-segments. Sub-segments that are less than 3-frames long were excluded from the turning angle analysis, which was done based on (Wortel et al., 2021).

The pitch length is the displacement associated with one complete helical turn, which was quantified as the straight-line distance between the starting and ending points of the turn. These points were identified through visual inspection of the trajectories and parasite orientation in 3-D space for the pitch calculation in Fig 4C.

The 3-D reconstruction of the processed DIC image stacks, along with the tracks exported from TrackMate (Fig 3C, Fig 4B, Video S1-S4), was done using the 3-D viewer and Track Processor in Icy (de Chaumont et al., 2012). Gaussian blur with a sigma of 2 was applied to the processed stacks before Icy rendering.

Outliers were removed from the datasets before performing statistical analysis. For each measured variable (Fig 4A&C, Fig 5), all the data points were combined to calculate the first (Q1), third (Q3) quartiles, and interquartile range (IQR). A data point is defined as an outlier if it is 1.5 times the IQR greater than the Q3 or 1.5 times the IQR less than the Q1.

The Kruskal-Wallis test was done to compare the groups for all of the track measurements in Figures 4A&C and Figure 5 (RStudioTeam, 2020). We used R version 4.1.2 (Team, 2021) and the following R packages: data.table v. 1.14.6 (Dowle and Srinivasan, 2022), gridExtra v. 2.3 (Auguie, 2017), onewaytests v. 2.7 (Dag et al., 2018), pracma v. 2.4.2 (Borchers, 2022), rmarkdown v. 2.20 (Allaire et al., 2023; Xie et al., 2018; Xie et al., 2020), rstatix v. 0.7.2 (Kassambara, 2023), scales v. 1.2.1 (Wickham and Seidel, 2022), tidyverse v. 2.0.0 (Wickham et al., 2019).

### MIC2 secretion assay

To examine microneme secretion, freshly egressed parasites were harvested. 4×10^7^ parasites were resuspended in 90 μl DMEM with 1% bovine calf serum, incubated at 10 min at 37 °C with or without 5 µM A23187, placed on ice for 5 min, and centrifuged for 10 min at ~ 800 g to separate the secreted fraction (supernatant) from the parasites (pellet). The supernatant (40 μl) and pellet fractions were analyzed by Western blot as described in (Leung et al., 2019) with a mouse 6D10 anti-TgMIC2 antibody (Carruthers et al., 2000) at 1:1000 dilution, a mouse anti-tubulin (Sigma, T6074) at 1:4000 dilution, and a Donkey anti-mouse IRDyE 680RD (Li-cor #926-68070) at 1:7000 as secondary antibody. To normalize A23187-induced MIC2 secretion, background-subtracted fluorescence of the MIC2 band from the supernatant sample was divided by background-subtracted fluorescence of the tubulin band in the corresponding pellet sample.

### Egress assay

For induced egress assay, parasites were added to MatTek dishes containing a confluent HFF or BS-C-1 monolayer and grown for ~ 35 hours. Cultures were washed once and incubated in 2 ml of L15 imaging media and placed at ~7°C for 1 hr, before being moved to the environmental chamber kept at 37°C. The media was then replaced with ~ 2 ml of 5 µM A23187 in L15 imaging media. Images were collected at 37°C with 1 sec interval for 10 minutes on an Applied Precision Delta Vision imaging station equipped with an environmental chamber.

### Invasion assays

To analyze the behavior of live parasites during invasion, extracellular parasites were used to infect a MatTek dish containing a nearly confluent monolayer of HFF cells. The dishes were incubated on ice in L15 imaging media for ~10 min to facilitate parasite settling on the host cells. The culture was then imaged at 37°C with 1 sec intervals for 15 min to capture invasion events.

Immunofluorescence-based invasion assays were carried out as described in (Munera Lopez et al., 2022) with some modifications. ~4×10^6^ freshly egressed parasites were used to inoculate a MatTek dish of nearly confluent HFF cells. After 10 min incubation on ice and then 10 min incubation at 37°C, the samples were processed for immunofluorescence. Three biological replicates, each with three technical replicates, were performed. Parasites in 15 fields were counted for each strain per technical replicate. In total (intracellular + extracellular), 4852 parasites were counted for the RH*Δhx* parent and 2944 for the TKO parasite. P-values were calculated by unpaired Student’s t-tests for the biological replicates.

### Plaque assays

A total of 50, 100 and 150 freshly harvested parasites were added to each well of 6-well plates containing confluent HFF monolayers and grown undisturbed for 8 days at 37°C, then fixed with 70% ethanol for 15 min and stained with 1% (wt/vol) crystal violet in 25% (vol/vol) methanol, rinsed with PBS, air-dried, and imaged.

## Acknowledgments

We are grateful to Dr. Owen Saxton (University of Cambridge, United Kingdom) for providing the source code for his ‘Semper’ image processing package, Dr. Con Beckers (William Carey University, Mississippi) for the rabbit anti-TgIMC1 and anti-TgGAP45 antibodies, Dr. Peter Bradley (University of California, Los Angeles) for the mouse anti-TgISP1 antibody, and Dr. Vern Carruthers (University of Michigan, Ann Arbor) for the mouse mAb 6D10 anti-TgMIC2 antibody. We thank Melissa Molina for tissue culture support.

## Supplemental Videos and Figures

**Video S1:** Time-lapse microscopy of 3-D motility in 50% Matrigel for the RH*Δhx* (RH parental 1) and TKO parasites. *Left:* Videos of projected DIC images (projections of 3-D stack generated by Stack focuser in ImageJ/Fiji). *Right:* Corresponding processed images with a subset of 3-D trajectories. Time interval: ~ 1.36 sec. Video speed: 10 frames/s.

**Video S2:** Rotational view of 3-D trajectories for the RH*Δhx* (RH parental 1) and TKO parasites included in Video S1. The traveling sphere associated with each track represents the centroid of the parasite traveling along that path. Length of arrow= 16 μm

**Video S3:** Time-lapse microscopy of 3-D motility in 50% Matrigel for the RH*Δku80* (RH parental 2) and *Δdcx* (*dcx KO*) parasites. *Left:* Videos of projected DIC images (projections of 3-D stack generated by Stack focuser in ImageJ/Fiji). *Right:* Corresponding processed images with a subset of 3-D trajectories. Time interval: ~ 1.36 sec. Video speed: 10 frames/s.

**Video S4:** Rotational view of 3-D trajectories for the RH*Δku80* (RH parental 2) and *Δdcx* (*dcx KO*) parasites included in Video S3. The traveling sphere associated with each track represents the centroid of the parasite traveling along that path. Length of arrow= 16 μm

**Video S5:** Time-lapse microscopy of A23187 induced-egress for RH*Δhx* (RH parental) and TKO parasites. A23187 was added at the beginning of the movies to a final concentration of 5 µM. The intracellular wild-type and TKO parasite cultures were placed at 7°C for 1 hr prior to the egress experiment, which was conducted at 37°C. Time interval: 1 sec. Video speed: 60 frames/s. Scale bar: 5 µm.

**Video S6:** Time-lapse microscopy of host cell entry for RH*Δhx* (RH parental 1), TKO and RH*Δku80* (RH parental 2) and *Δdcx* (*dcx KO*) parasites. Fig 6F includes images selected from the timelapses in the leftmost column. Time interval: 1 sec. Video speed: 6 frames/s. Scale bar: 5 µm.

**Figure S1.**
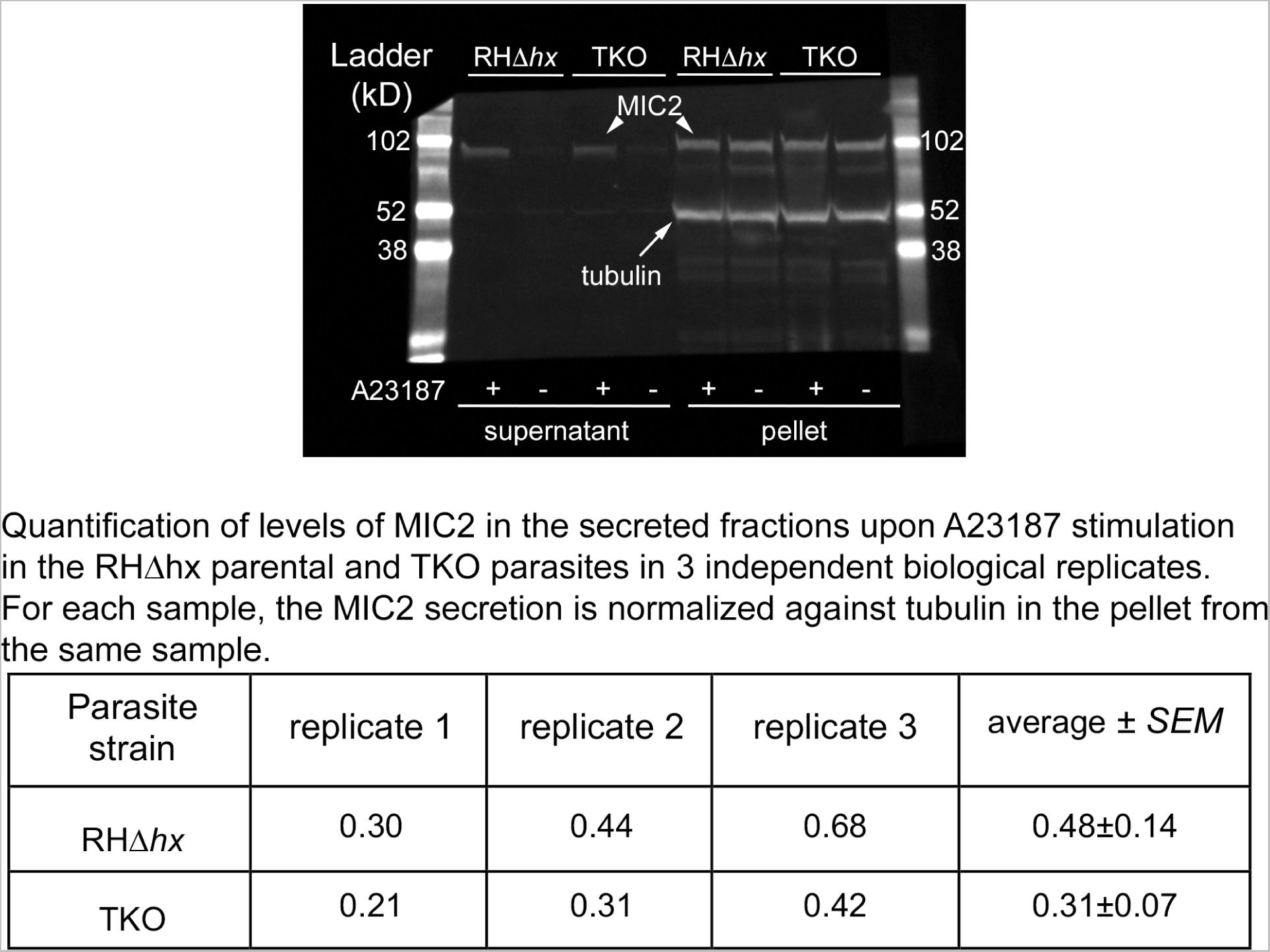
(Blot Transparency) *Top:* Image of Western blot that contains data displayed in Fig 6C. Western blot of the secreted (supernatant) and unsecreted (pellet) fractions of RH*Δhx* and TKO parasites with (+) or without (-) A23187 treatment. The blot was probed by antibodies against MIC2 and tubulin. Lanes for molecular weight ladders are included. *Bottom:* Quantification of levels of MIC2 in the secreted fractions upon A23187 stimulation in the RH*Δhx* parental and TKO parasites in 3 independent biological replicates.

**Figure S2.**
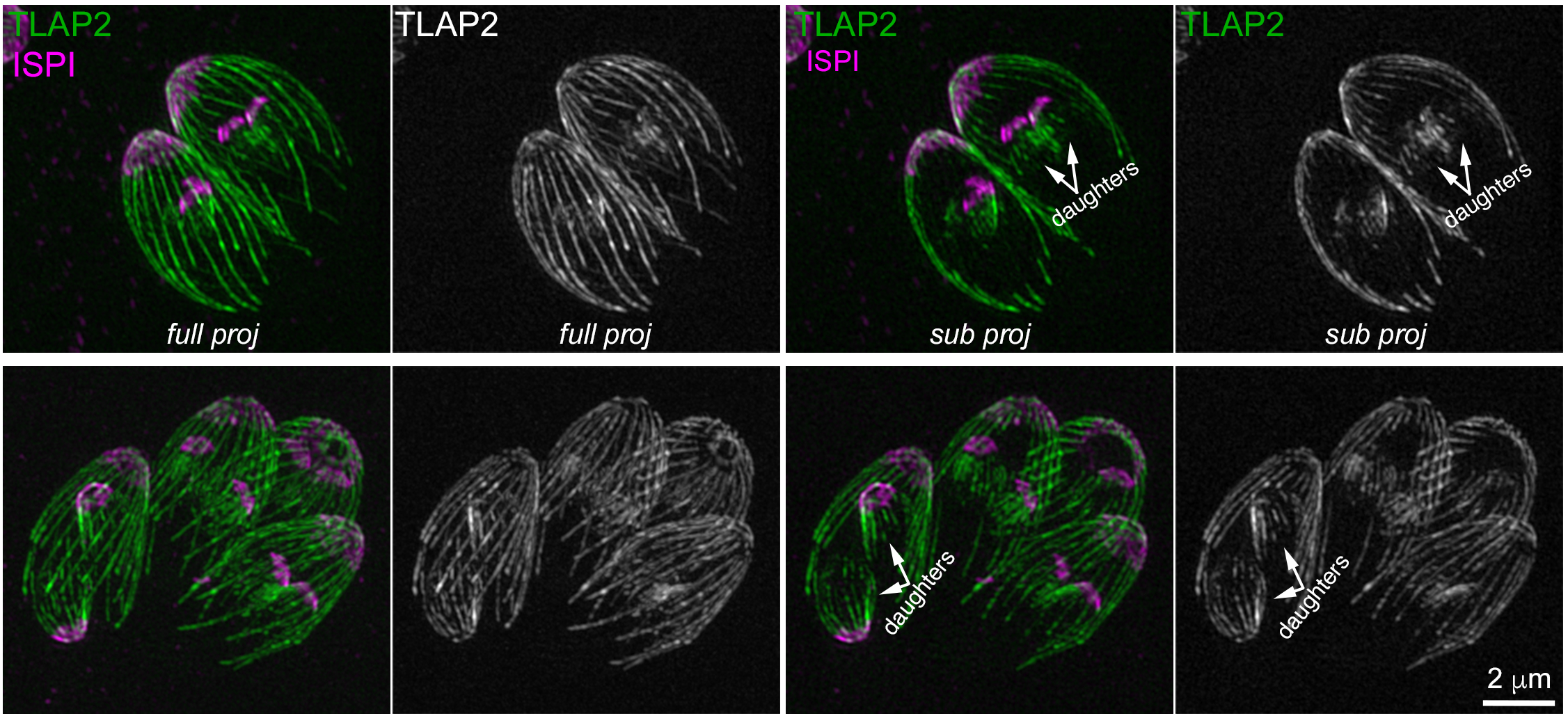
Full and subset projections of 3D-SIM image stacks of dividing intracellular *mEmeraldFP-TLAP2* knock-in parasites showing that TLAP2 is recruited to the daughter cortical microtubules. Green and grayscale: mEmerald-FP-TLAP2. Magenta: apical cortex labeling with a mouse anti-ISP1 antibody. Parasites at an earlier (top) and a later stage (bottom) of daughter formation are shown.

